# Inter-kingdom microbial interactions revealed by a comparative machine-learning guided multi-omics analysis of industrial-scale biogas plants

**DOI:** 10.1101/2023.02.07.527480

**Authors:** Roland Wirth, Zoltán Bagi, Prateek Shetty, Márk Szuhaj, Teur Teur Sally Cheung, Kornél L. Kovács, Gergely Maróti

## Abstract

Multi-omics analysis is a powerful tool for the detection and study of inter-kingdom interactions, such as those between bacterial and archaeal members of complex biogas-producing microbial communities. In the present study, the microbiomes of three industrial-scale biogas digesters, each fed with different substrates, were analysed using a machine-learning guided genome-centric metagenomics framework complemented with metatranscriptome data. This data permitted us to elucidate the relationship between abundant core methanogenic communities and their syntrophic bacterial partners. In total, we detected 297 high-quality, non-redundant metagenome-assembled genomes (nrMAGs). Moreover, the assembled 16S rRNA gene profiles of these nrMAGs showed that the phylum Firmicutes possessed the highest copy number, while the representatives of the Archaeal domain had the lowest. Further investigation of the three anaerobic microbial communities showed characteristic alterations over time but remained specific to each industrial-scale biogas plant. The relative abundance of various microbes as revealed by metagenome data were independent from corresponding metatranscriptome activity data. Interestingly, Archaea showed considerably higher activity than was expected from their abundance. We detected 53 nrMAGs that were present in all three biogas plant microbiomes with different abundances. The core microbiome correlated with the main chemical fermentation parameters and no individual parameter emerged as a predominant shaper of community composition. Various interspecies H_2_/electron transfer mechanisms were assigned to hydrogenotrophic methanogens in the biogas plants that ran on agricultural biomass and wastewater. Analysis of metatranscriptome data revealed that methanogenesis pathways were the most active of all main metabolic pathways. These findings highlight the importance of a combinatorial omics data framework to identify and characterise the activity of specific microbes in complex environments.

## 1. Introduction

Engineered anaerobic digestion (AD) systems produce biogas by efficiently recycling nutrients via the decomposition and utilisation of organic compounds [1, 2]. This process involves a diverse microbial community, with each microbe having a specific role. Previous research suggests that the assembly of AD microbial communities is not random, but is determined by factors such as microbe-microbe interactions, substrate composition and operational conditions [3–5].

Unravelling microbial interactions and their underlying mechanisms is an intricate task, since many microbes cannot survive without specific microbial partners [6]. Furthermore, over 99% of prokaryotes cannot be traditionally cultivated under laboratory conditions [7, 8]. The strong syntrophic relationships between microorganisms in AD microbial consortia therefore necessitate extensive study to determine their diversity, metabolic function and distribution. These studies have been initially hindered by limitations inherent to culture-dependent microbiological methods, which require the isolation of microorganisms and can be challenging when syntrophic relationships are ubiquitous [9–11]. High-throughput sequencing and bioinformatics tools permit bulk analysis of genomic material and thereby provide insight into the taxonomy and functions of entire microbial communities [12–14]. Biogas-producing communities contain a large and diverse contingent of uncharacterised microbes that have been described as a ‘microbial black box’ [15–17]. To address this issue, increasingly sophisticated bioinformatic algorithms can be used to reconstruct the genomes of individual species (or MAGs: metagenome-assembled genomes) from these mixed communities. A framework that combines a genome-centric metagenomics approach with metatranscriptomics analyses can be a powerful tool for understanding the taxonomic structures and metabolic capabilities of AD microbial communities without relying on reference-based techniques. More importantly, this framework would also permit the discovery of novel taxonomic groups and active functions [6, 16, 18].

Various strategies and algorithms have been proposed to recruit MAGs. [19]. Conventional reference-independent methods generally rely on sequence features (e.g. k-mer frequencies) and co-abundance statistics (e.g. coverage of reads mapped to individual contigs), which should vary more between genomes than within them. Several recent studies employing genome-resolved metagenomics have recovered MAGs from lab- and industrial-scale biogas reactors [20–23]. One recently published study showed that the use of a semi-supervised machine learning approach could significantly improve binning performance [24, 25]. For this approach, labelled data (‘cannot-link’) and unlabelled data (‘must-link’) are used to identify the feature distribution - i.e. whether certain pairs of contigs should be grouped together. The resulting distribution shape (‘info map’) generated by this binning process can then be integrated with contig characteristics (‘weighted k-means’) using machine learning to obtain high-quality MAGs.

In AD microbiomes, the identification of microbes common to many biogas plants can reveal major pathways and ecological features involved in the microbial food chain [26, 27]. Previous studies using 16S rRNA gene amplicon sequencing showed that core microbes create a stable community capable of resisting various perturbations (i.e. AD parameter changes) [15, 28, 29]. However, amplicon sequencing and read-based metagenomics approaches may be unable to identify unknown species by core microbiome analysis due to its dependence on reference databases [21, 30]. As a result, new bioinformatic protocols must be developed to discover the structure and function of unknown microorganisms [31, 32].

The present study investigated the structure and function of microbiomes from three industrial-sized biogas reactors, each fed with a distinct, characteristic substrate. Each reactor was monitored at seasonal intervals over a one-year period. Shotgun sequencing and a machine-learning-guided genome-centric metagenomic and metatranscriptomic analysis framework were used to identify microbial composition and ongoing metabolic activities. Next, the shared portion of the microbial community involved in digesting heterogeneous substrates was investigated using an occurrence-based core community concept [33]. The main finding of this study was the elucidation of a partnership between the abundant core methanogenic community and specific syntrophic microbes. More specifically, we describe key networks of interspecies syntrophy and find an intriguing correlation between chemical fermentation parameters and the core anaerobic microbiota present in the reactors.

## 2. Materials and Methods

### 2.1 AD samples

Samples were taken from three state-of-the-art anaerobic digesters in Hungary. Two of the digesters (MWBP and SZBP) are in Szeged and the other is in Kecskemét (KBP). Key characteristics of each of the biogas plants studied here are summarised in Supplementary Table 1. These biogas plants were selected using the following criteria: (i) the digesters have been operating without issue for more than five years and (ii) the digesters use distinct biopolymers as the main substrate of decomposition for biogas production. Sampling was performed for a one-year period at the following (seasonal) intervals: Oct 2020, Jan 2021, Apr 2021 and Jul 2021. Samples were directly transported to the laboratory and were processed immediately on arrival. AD parameter measurements and DNA/RNA purification were performed on fresh biogas plant (BP) samples in triplicate (i.e. biological triplicates: n = 3).

### 2.2 Determination of AD chemical parameters

For each BP, we measured: sludge pH, carbon to nitrogen ratio (C/N), total solid (TS) content, volatile solid (VS) content, total ammonia nitrogen (TAN; i.e. ammonium ions and dissolved ammonia), volatile organic acid (VOA) content and total inorganic carbon (TIC). Measurements were taken as previously described in Wirth *et al.* 2021 [34].

Biochemical methane potential (BMP) batch tests were performed in 160 mL reactor vessels (Wheaton glass serum bottle, Z114014 Sigma-Aldrich) containing 60 mL liquid phase. The substrate (α-cellulose: C8002 Sigma-Aldrich) to inoculum ratio was set according to the VDI 4630 standard (substrate to inoculum ratio = 2:1 g VS; Vereins Deutscher Ingenieure 4630, 2006). Filtered inoculum sludge (i.e. after removal of particles larger than 2 mm) was used for BMP measurement. A detailed description of the gas sampling and measurement procedures can be found in Szilágyi *et al.* 2021 [35].

### 2.3 Total DNA and RNA purification

Aliquots of 2 mL were obtained from the samples of each BP for total community DNA and RNA isolation. Purification was performed in triplicate and the resulting extractions were pooled together. All extractions were carried out using ZymoBIOMICS DNA/RNA miniprep kits (R2002, Zymo Research, Irvine, USA). After lysis (bead homogenisation was performed using a Vortex Genie 2 with a bead size of 0.1 mm, a homogenisation time of 15 min and at max speed), the Zymo Research kit parallel DNA and RNA purification protocol was followed. DNA and RNA quantity was estimated using an Agilent 2200 TapeStation (Agilent Technologies, Santa Clara, USA).

### 2.4 Metagenome and metatranscriptome sequencing

We closely followed all manufacturer recommendations for the Illumina sequencing platform (Illumina Inc., San Diego, USA). Pooled genomic DNA samples were used to sequence libraries constructed using the NEBNext Ultra II Library Prep Kit (NEB, Ipswich, USA). Paired-end metagenomics sequencing was performed on an Illumina NextSeq 550 sequencer using NextSeq High Output Kit v2 sequencing reagent kit. Metatranscriptome sequencing from pooled RNA was performed as follows: paired-end libraries were first prepared using a Zymo-Seq RiboFree™ Total RNA Library Kit, which includes a universal rRNA depletion step. mRNA sequencing was then performed on an Illumina NextSeq 550 sequencer using the NextSeq High Output Kit v2 sequencing reagent kit. Primary data analysis (i.e. base-calling) was performed using ‘bcl2fastq’ software (version 2.17.1.14, Illumina). Characteristic fragment parameters are summarised in Supplementary Table 2.

### 2.5 Metagenome assembly and binning

Raw sequences were filtered by fastp (version 0.23.2, length required: 150bp) and checked with FastQC (version 0.11.8). The filtered sequences produced by fastp were then co-assembled separately by Megahit (version 1.2.9, 4 samples per BP = 3 co-assemblies). The settings used were as follows: min contig length = 1500; min k-mer size = 21; max k-mer size = 141 [36]. The metagenomics binning procedure was performed separately for each AD metagenomics dataset until the dereplication step. We used Anvi’o (version 7: ‘hope’) to create the contig database for the following metagenomics workflow [37].

Genome reconstruction was performed using Semibin (version 1.1.1), a machine-learning-guided software package that combines a semi-supervised approach with deep Siamese neural networks by using an advanced co-assembly binning workflow with a semi-supervised mode [24]. For dereplication and quality filtration of metagenome-assembled genomes (MAGs) we used dRep (version 2.2.3) and CheckM (version 1.1.6) with the following parameters: dereplicate: comp 10, con 5, S_algorithm fastANI, sa 0.95 [38, 39]. MarkerMAG was used in default mode to detect, assemble and link 16S rRNA genes to MAGs and calculate the corresponding copy number [40].

Open reading frames (ORFs) were identified by Prodigal (version 2.6.3). InterProScan version 5.31–70 was used to functionally annotate gene coding sequences using the Pfam database [41]. Functional profiles were supplemented using data from the Kyoto Encyclopedia of Genes and Genomes (KEGG) function modules by Anvi’o (anvi-run-kegg-kofams) [42]. The enzymes involved in carbohydrate utilisation were identified using a combination of Pfam functional profiles and data from the carbohydrate-active enzyme database (CAZy). We then used the Genome Taxonomic Database (GTDB: Release 207) with GTDBTk (version 2.1.1) for taxonomic assignment [43]. Reconstructed non-redundant MAGs (nrMAGs) that showed greater than 90% completeness were also compared with entries in the Biogas Microbiome database [20] using fastANI (version 1.33) [44].

nrMAG statistics are summarised in Supplementary Table 3. CoverM (version 0.6.1, https://github.com/wwood/CoverM; method: rpkm) was used to calculate the reads per kilobase per million mapped reads (RPKM) of the nrMAGs. Phylogenomic trees were generated using a set of 120 bacterial and 53 archaeal Single-copy Core Genes (SCGs) via GTDB-Tk (classify_wf) and IQTree (version 2.2.0.3) using the following parameters: number of bootstraps: 1000, maximum iteration: 1000, stopping rule: 100. The interactive Tree of Life (iTOL: version 6; https://itol.embl.de/) tool was employed to visualise the phylogenomic tree as well as some binning results.

### 2.6 Metatranscriptome mapping and analysis

We first extracted ORFs from MAGs using bedtools: getfasta to obtain MAG-specific gene calls. We then combined these to create a Salmon index (using the parameter: keepDuplicates; https://github.com/COMBINE-lab/salmon). Read counts in transcripts per million (TPM) were generated for each sample using Salmon (version 1.8.0) in quasi-mapping mode with GC bias correction (gcBias).

### 2.7 Statistical analysis

BMP test results were visualised by the *ggplot2* and significant differences between maximum biogas potentials (i.e. from batch tests using α-cellulose) were then calculated using the *ggsignif* package (i.e. using a between-pairs Wilcoxon-test, with one-way ANOVAs used to compare multiple groups). Euclidean distances, multidimensional scaling of samples (i.e. MDS plots to display Bray-Curtis dissimilarities), MAG RPKM values and TPM values were visualised by microViz [45]. Differences among and within samples were calculated using permutational multivariate analysis of variance (PERMANOVA: n_perms: 10000). To assess the significance of differences between MAGs, we used Benjamini-Hochberg-adjusted p-values (hereafter termed ‘padj’), with a significance threshold of p ≤0.05.

We then analysed occurrence data to identify the core microbiome. A taxon observed in 83.3% of samples (i.e. in 10 out of 12, to avoid missing rare microbes) was considered to be a member of the core microbiome [33]. Pearson correlations between fermentation parameters and taxonomic ID of core microbiome members (at the species level or otherwise at the highest taxonomic level) were calculated by microViz. The *ggtern* package was used to visualise the core microbiome (at the phylum level) distribution between the BPs.

The results of the metatranscriptomics analysis of carbohydrate-active enzymes was visualised by Circos online (http://circos.ca/) and *ggplot2* (version 4.1.3). To process metatranscriptomic data and calculate the log2 fold change (log2 FC) between the KEGG modules of different biogas plants, we used the *DESeq2* package implemented in R [46]. Finally, to visualise significantly different KEGG pathways we used the EnhancedVolcano package (https://github.com/kevinblighe/EnhancedVolcano) to display *DESeq2* results (using the following settings: pCutoff: 0.05, FCcutoff: 2.0).

## 3. Results and Discussion

### 3.1 Different substrates but similar methane yields

Samples were taken from the anaerobic digesters of three industrial-scale biogas plants (further information: section 2.1 and Supplementary Table 1). KBP was mainly fed with chicken manure and pretreated wheat straw, SZBP was fed pig slurry and maize silage and MWBP was fed a highly diverse set of substrates, including municipal wastewater sludge (Suppl. Table 1). All digesters were operated at mesophilic (i.e. 36°C to 38°C) temperatures in continuously stirred tank reactors while under operation. During the seasonal monitoring period (i.e. four sampling points per biogas plant), all reactors operated stably and no failures were reported.

Standard BMP tests were performed (see Materials and Methods) to measure the maximum biogas potential of the different AD communities. Methane yields varied from 310 mL (Standard Deviation: 2.6 mL) to 376 mL g VS^−1^ (SD: 14.6 mL). Although the inoculum for BMP tests originated from digesters fed with distinct substrates, the methane yields were highly similar (Fig. 1A). Moreover, tests from the Oct, Jan, Apr and Jul timepoints all showed similar methane yields among the three biogas plants (Fig. 1B; For April: KBP = 362 ± 14.6; MWBP = 356 ± 2.4 and SZBP=365 ± 7.6 mL methane g VS^−1^). In previous studies, the ADs of two different agricultural waste and wastewater sludge sources also showed similar methane potential ranges [47–49].

**Figure 1:**
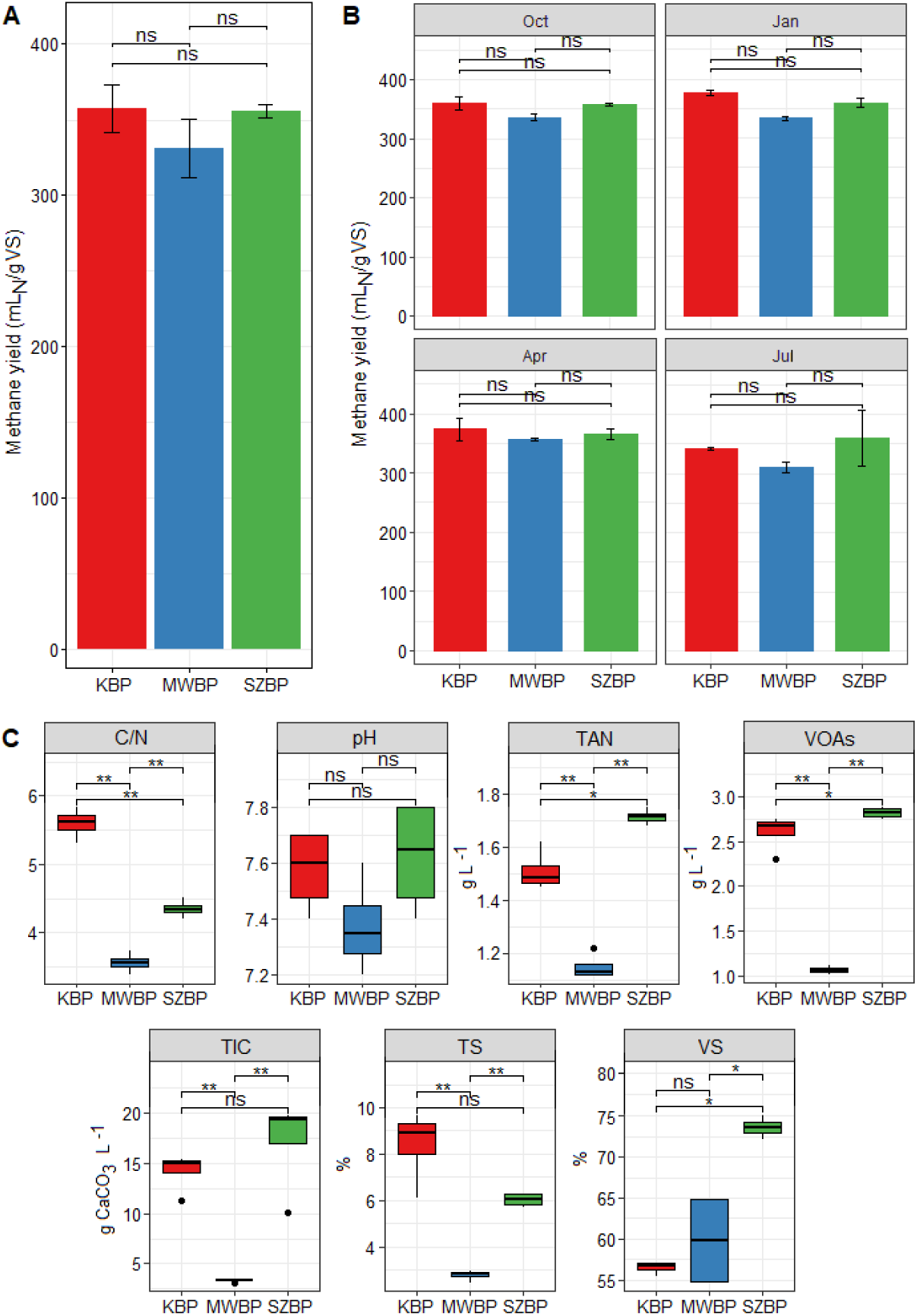
Parameter measurements of the AD chemical process. Different colours represent the three BPs. The Kecskemét biogas plant (KBP) is shown in red, the biogas plant digesting municipal wastewater sludge (MWBP) is shown in blue and the Szeged biogas plant (SZBP) is shown in green. A BMP methane yields of the three biogas plant substrate mixes (see section 2.1). B BMP methane yields over the tested periods. C Anaerobic digestion chemical parameters of the sludge from individual BP digesters (VOAs: volatile organic acids, TIC: total inorganic carbon, TAN: total ammonia nitrogen, TS: total solids, VS: volatile solids, C/N: carbon to nitrogen ratio). Mean differences were analysed via ANOVA and considered statistically significant as follows: p < 0.05 (*), p < 0.001 (**).

Although the main chemical process parameters were within optimum ranges [50–52], they showed clear differences among the three BP reactors (Suppl. Table 1). We observed the following range of values for different parameters as follows: volatile organic acids (VOAs) [1 g L^−1^ (MWBP) to 2.9 g L^−1^ (SZBP)]; total inorganic carbon (TIC) [3.1 g CaCO_3_ L^−1^ (MWBP) to 19.8 g CaCO_3_ L^−1^ (SZBP)]; total ammonia nitrogen (TAN) [1.1 g L^−1^ (MWBP) to 1.7 g L^−1^ (SZBP)]; carbon to nitrogen ratio (C/N) [3.3 (MWBP) to 5.8 (KBP)]; total solids (TS) [2.4% (MWBP) to 9.6% (KBP)] and volatile solids (VS) [54.7% (MWBP) to 75.1% (SZBP)] (Fig. 1C). Of these, the C/N, VOAs and TAN concentrations showed the highest variability among the BPs (ANOVA: p < 0.05). Moreover, these values were typically lower in MWBP (p < 0.01) [53].

### 3.2 High-quality MAGs from the AD microbiomes

Semi-supervised binning complemented with machine learning (ML) is a newly developed approach that has proven to be highly beneficial for developing knowledge of microbial genomics. Pan *et al.* (2022) found that a habitat-specific model originated from ML improved the metagenome binning process for complex microbiomes; it outperformed extant unsupervised, nucleotide composition and abundance-based methods (i.e. Metabat2) [24, 25]. However, due to unusual read-depth and atypical nucleotide composition, ribosomal RNA genes (especially 16S) are frequently absent from MAGs recovered from short-read sequencing data [32, 40]. These genes are essential for studying the genomics and phylogeny of uncultivated microbes and permit analyses connecting MAG data with 16S rRNA gene databases.

In the present study, more than 296 million metagenomic sequence reads passed the filtering step (with an average 24.6 million reads per sample). Filtered reads were co-assembled by Megahit (three independent assemblies per BP), resulting in a total of 283,491 contigs (KBP: 107,920; MWBP: 98,415; SZBP: 77,156 contigs; for details see: Suppl. Table 2). A genome-centric metagenomics strategy was followed for each co-assembled contig set. The dereplicated and quality-filtered set of MAGs was then used for further analysis. Semibin produced 297 nrMAGs (nr = non-redundant) of which 107 (36%) had more than 90% completeness. It is noteworthy that 27% of nrMAGs were identified as containing 0% contamination by CheckM analysis (Fig. 2A and B). Furthermore, 24% of nrMAGs (n = 70) were identified at a quality level that was higher than their representatives in the Genome Taxonomy Database (Suppl. Table 3). These observations confirm the effectiveness of Semibin as a binning procedure for complex anaerobic biogas-producing communities [24]. In addition, the larger number of unique nrMAGs binned by Semibin may also lead to better mRNA mapping.

**Figure 2:**
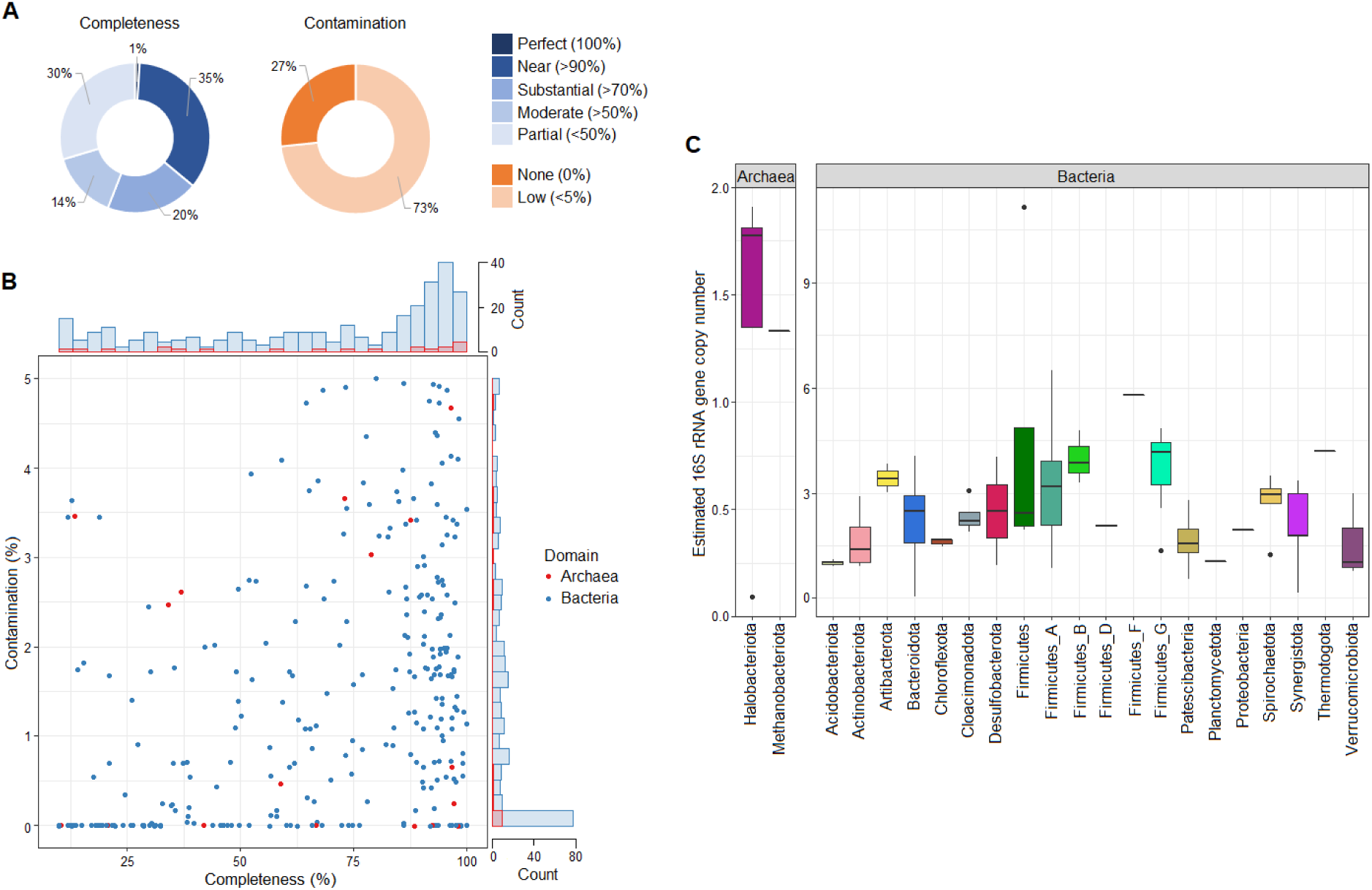
Binning performance and 16S rRNA gene copy number estimation of metagenomics data originated from complex biogas-producing microbiomes. **A** Comparison of the completeness and contamination of the non-redundant metagenome-assembled genomes (nrMAGs) produced by Semibin and analysed by CheckM using the default set of SCGs. **B** Distribution of nrMAGs reconstructed by Semibin based on completeness and contamination. **C** Estimated 16S rRNA gene copy number for 22 phyla (i.e. two Archaea and 20 Bacteria taxa).

The MarkerMAG programme was then used to detect, assemble and link 16S rRNA genes from metagenomes to MAGs and to estimate copy numbers (see: Materials and Methods section 2.5 and Suppl. Table 3) [40]. In the present study, 16S rRNA genes (min length: 1200 nucleotides) were detected in 82 nrMAGs distributed across 22 phyla (representing 28% of all nrMAGs). Representatives of the Firmicutes possessed the highest estimated mean copy number of this gene (3.6 copies), while members of the Halobacteriota and Methanobacteriota showed the lowest copy numbers, with an average of 1.3 copies each (Fig. 2D). Along with previous scientific reports, our data suggested that amplicon sequencing of 16S rRNA underestimated the abundance of the archaeal community in AD [54, 55]. One method to better estimate microbial abundance derived from 16S rRNA gene sequencing is to normalise results via copy number per detected genome [56]. However, the actual 16S rRNA gene copy number is unknown for many prokaryotes [32]. Current bioinformatic solutions for normalising amplicon sequencing data rely on the apparent phylogenetic conservation of individual gene copies and this assumption may only be valid for short phylogenetic distances [57]. Genome-resolved metagenomics combined with 16S rRNA gene detection may help to bridge this knowledge gap.

### 3.3 BPs showed dissimilar microbiome compositions

Overall, we found that the three AD reactors harboured distinct microbial communities. Multidimensional scaling (MDS; implemented using the Bray-Curtis dissimilarity measure) revealed that the variance in microbiome composition across the three biogas plants was significantly larger (PERMANOVA: p = 0.001) than the variance in microbiome structure across different sampling points of each individual biogas plant (PERMANOVA: p = 0.34). The microbiomes of the different BPs showed characteristic changes over time, but in each case these changes were distinctive for that plant (Fig. 3B). This finding was confirmed by Euclidean distance calculation, which indicated that the microbiomes of KBP and SZBP were more similar to each other than to MWBP (Fig. 3A). In accordance with past research, our analyses confirmed that the KBP and SZBP microbiomes predominantly contained Bacteroidia, Clostridia and Limnochordia, whereas the MWBP microbiome contained Bacteroidia, Anaerolineae and Actinomycetia (Fig. 3 D) [20, 23, 58]. We can therefore conclude that there is a common microbial community landscape that includes the main taxa found in the biogas digesters studied here [3, 4].

**Figure 3:**
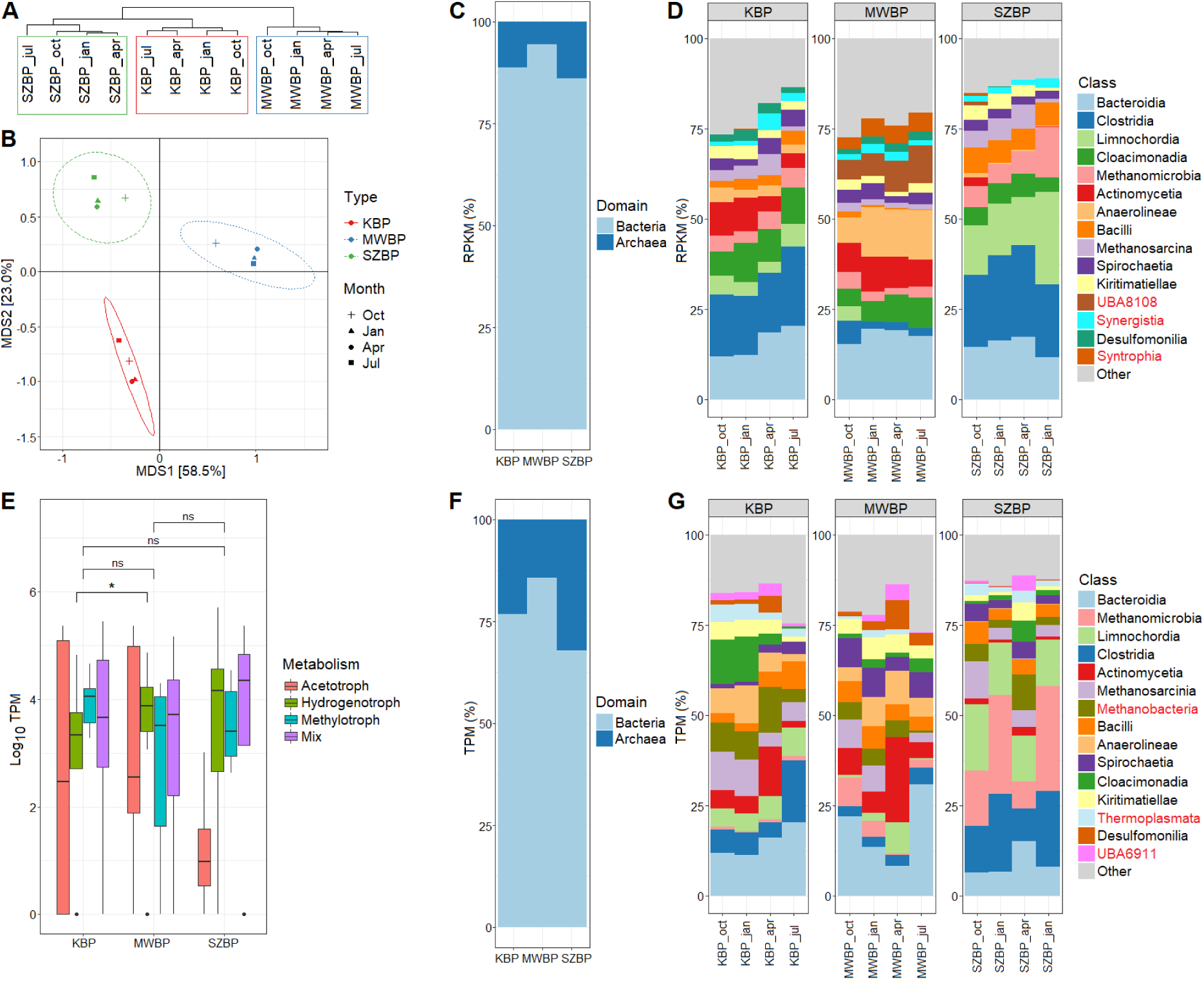
Biogas-producing microbiome abundance (in RPKM %) and activity (in TPM %) analysed using a genome-centric metagenomics and metatranscriptomics framework. **A** Euclidian distance of BP samples. This measure represents the dissimilarity of microbial communities at the level of species annotation or higher. **B** Multidimensional scaling of samples. This measure indicates the Bray-Curtis dissimilarities of the microbial community at the level of species annotation or higher. Each symbol is related to a specific BP sample. Oct, Jan, Apr and Jul, indicate the months in which the samples were collected. **C** Taxonomic distribution of nrMAGs at the level of domain for the three BPs (in RPKM %). **D** Taxonomic distribution of nrMAGs. This shows the most abundant taxa in all twelve samples taken (i.e. each BP-time point pair) resolved to the class taxonomic level (shown in RPKM %). **E** The overall metatranscriptomic activity of archaeal microbes involved in different methanogenic pathways (mix: indicates those nrMAGs that are capable of using all three methanogenic pathways). **F** Cumulative metatranscriptomic activity of nrMAGs at the domain taxonomic level for the three BPs (shown in TPM %). **G** The metatranscriptomic activity of nrMAGs. Shown are the most active taxa present in all twelve samples taken (i.e. each BP-time point pair) resolved to the class taxonomic level (shown in TPM %).

Examination of microbial abundance and activity using metagenome and metatranscriptome data distributions revealed distinct patterns. Based on relative abundance (RPKM %), the top three classes found were Bacteroidia, Clostridia, and Limnochordia, although the activity (TPM %) of Methanomicrobia outperformed members of these classes. Moreover, we also found striking differences in rank and in taxonomic composition of the top 15 most active microbes compared to the most abundant microbial classes (Fig. 3D and G). Thus, the relative abundance of microbes is divergent from relative metatranscriptomic activity. Overall, the representatives of the domain Archaea showed higher activities than abundance, as evidenced by comparing TPM % to RPKM % values (Fig. 3C and F; Suppl. Tables 4 and 5). This phenomenon was also observed in an anaerobic biogas-producing community analysed by De Vrieze *et al.* (2016) [59].

Interestingly, in all three industrial-scale biogas plants, the methanogenic Archaea identified using metatranscriptomics showed no significant differences in overall activity (Fig. 3E). This implies the functional stability of the archaeal community using multiple pathways for methane production [15, 60]. Only hydrogenotrophic methanogens, which use CO_2_ and H_2_ as substrates for methane generation, exhibited different activities in KBP and MWBP (Wilcoxon-test: p < 0.05), indicating the presence of highly active and diverse methanogenic functions in MWBP.

### 3.4 Biogas-producing communities and the ‘microbial black box’

Based on known lineage-specific marker gene sets (SCGs) and on MIMAG data, we detected 36% high-, 34% medium- and 30% low-quality nrMAGs [61]. The Genome Taxonomy Database (GTDB) was then employed for the taxonomic assignment of reconstructed nrMAGs. These results showed that nrMAGs could be categorised into 32 phyla, from which three belonged to the Archaea and 29 belonged to the bacteria (Fig. 4 and Suppl. Table 3).

**Figure 4:**
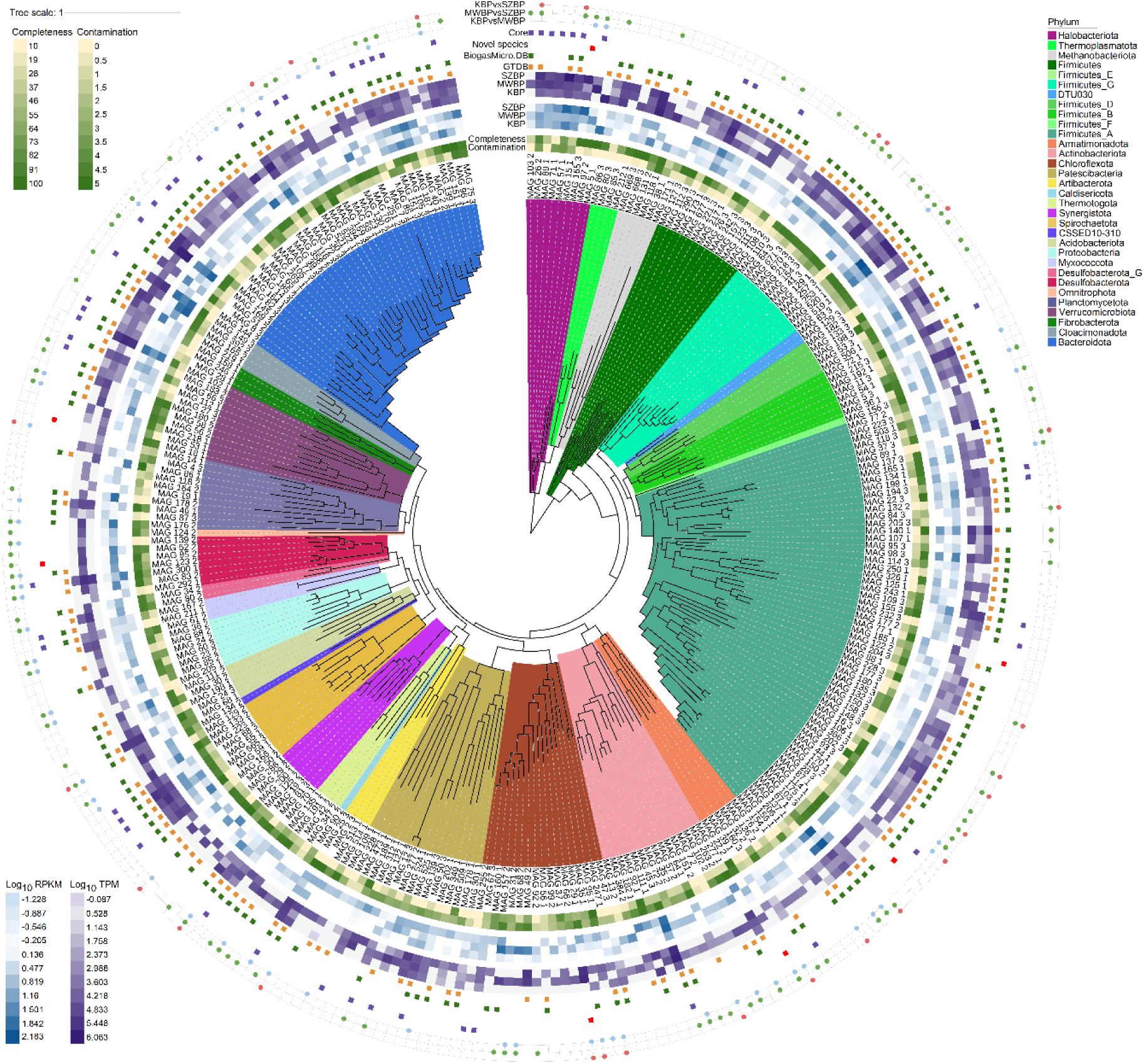
Phylogenetic tree of reconstructed nrMAGs based on bacterial and archaeal SCGs. The background colour of the inner phylogenetic tree marks the phylum that they belong to. The first ring shows the nrMAG number. The next section contains the quality metrics of nrMAGs. The second and third heatmap rings represent the cumulative abundances (in log10 RPKM) and metatranscriptomic activity (in log10 TPM) for the nrMAGs sourced from KBP, MWBP and SZBP, respectively. In the next two rings the symbols represent the presence of specific nrMAGs in the GTDB (orange) and Biogas Microbiome (green) databases. The red symbols mark the seven nrMAGs that have high completeness (>90%), low contamination (<5%) and have been not found in any available databases (Archaea: MAG_165_3; Bacteria: MAG_114_1, _18_1, 29_1, 77_1, 87_1 and 95_2. Purple symbols represent the core nrMAGs. The outer ring shows the significant differences using Benjamini-Hochberg-adjusted p-values (p < 0.05).

The nrMAGs were also compared to the Biogas Microbiome database by using average nucleotide identity (ANI) calculation implemented by fastANI [20]. This database contains a comprehensive set of microbial genomes previously found in biogas digesters. We found that 83% of the reconstructed high-quality nrMAGs (completeness >90%; n = 107) were present in the Biogas Microbiome database (ANI cutoff: 95%). Interestingly, seven high-quality nrMAGs, representing 7% of the high-quality nrMAGs (n = 107) and 2% of all nrMAGs (n = 297) were not associated with a closest representative in either database; that is, they did not meet the following thresholds: GTDB: species-level identification with 95% reference radius and Biogas Microbiome: ≥ 95% ANI (Fig. 4 and Suppl. Table 3). Notably, three high-quality nrMAGs were found to contain 16s rRNA sequences as well (Suppl. Table 3). The seven putative novel nrMAGs also showed both high abundance and activity, at 3% RPKM and 5% TPM with respect to the whole analysed microbiome, respectively (Suppl. Table 3 and 4). These nrMAGs may be members of the hypothesised ‘microbial black box’ that can now be released into the realm of known participants in AD communities.

### 3.5 AD chemical parameters and the core community

Genome-centric metagenomics-based core microbiome analysis was used to identify potentially relevant taxa for the shaping of the core microbial community. This macroecological study was therefore supported by metatranscriptome data (Fig. 5 and Suppl. Figure 1) [33]. These analyses provide insight into the relationship between nrMAGs and AD parameters, as well as their roles in shaping the core microbiome.

**Figure 5:**
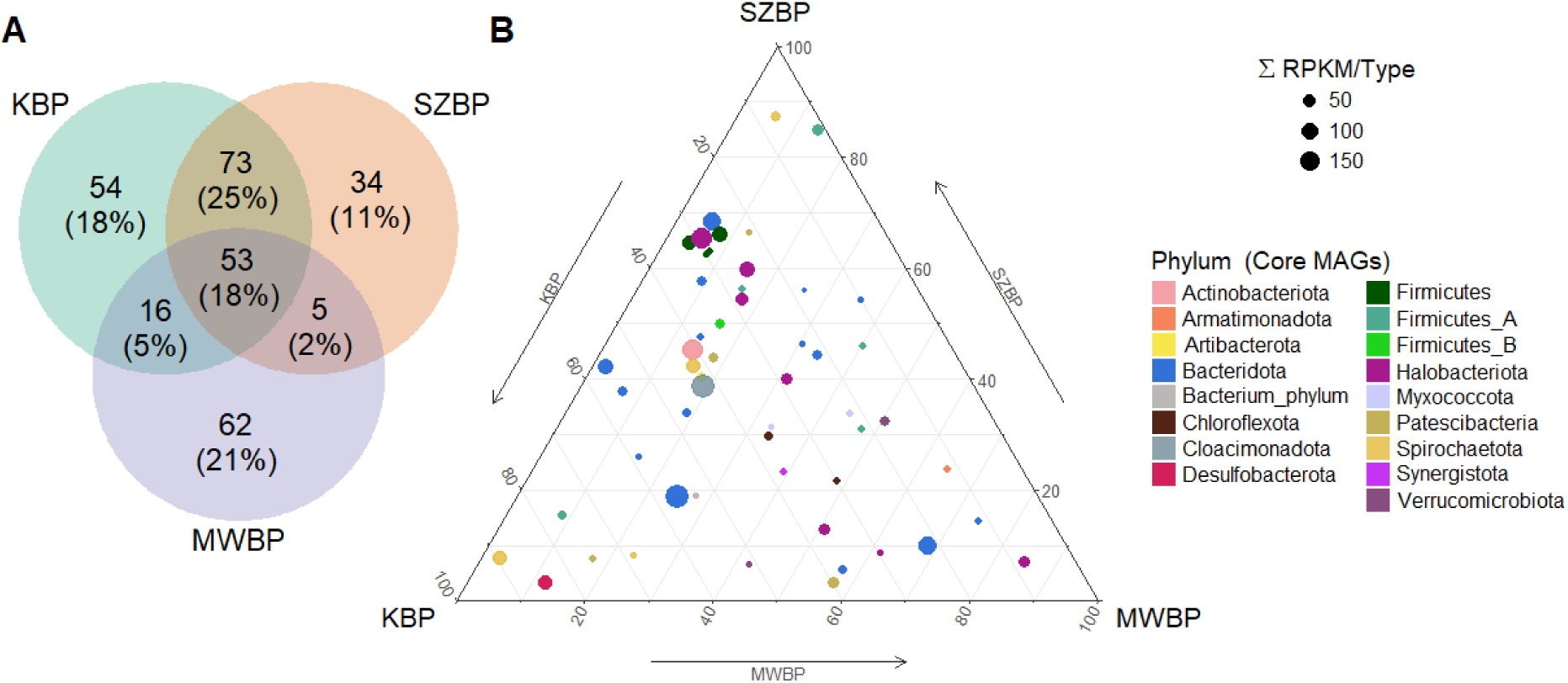
Analysis of core microbiome distribution. **A** Venn diagram indicates the total number of nrMAGs and the number of shared nrMAGs among the three BPs (percentages as indicated). **B** Ternary plot showing the distribution of bacterial and archaeal nrMAGs between BPs. Dot colour represents the phylum that the nrMAGs belongs to. The size of the dot is proportional to total abundance (i.e. cumulative RPKM value in all BPs) of specific nrMAGs.

The examination of the distribution of reconstructed nrMAGs in specific biogas plants revealed that MWBP possessed the most unique microbiome of the three investigated systems. KBP and SZBP showed somewhat overlapping microbiomes, as evidenced by the MDS and Euclidean distance calculations. Nonetheless, we detected 53 nrMAGs in all three digesters (Fig. 5A). Most of the core microbes identified were representatives of well-known hydrolysing bacteria (e.g. Firmicutes, Bacteriodota, etc.) and hydrogenotrophic methanogens (e.g. Halobacteriota). These microbes play an important role in maintaining biogas productivity and sustainable system performance [12, 62]. Although some nrMAGs were present in all digesters, their abundances varied considerably. Among core bacteria, the phyla Firmicutes and Bacteriodota were found in both SZBP and KBP, while members of the Verrucomicrobiota and Armatimonadota predominated in MWBP [20, 53] (Fig. 5B).

Of the core nrMAGs, we selected the top 30 and correlated their abundance with each other and with AD chemical process parameters using Pearson correlations (See: Section 2.7) (Fig. 6). The main parameters influencing the abundance of top core microbes were TAN, VOAs and TIC, all of which showed Pearson correlations greater than 0.6. Based on this observation, five clusters showing characteristic correlations with AD chemical parameters were plotted (Fig. 6B). The microbes within the same cluster were positively correlated with one another, while the microbes in Clusters I, IV and V were negatively correlated with one another (Fig. 6A). We also found that top core nrMAGs belonging to the phyla Firmicutes, Spirochaetota and Caldatribacteriota were positively correlated with VS, TAN, VOAs and TIC, while members of the phylum Bacteroidota were also correlated with many of the measured parameters. Finally, the C/N ratio showed a more pronounced impact on Bacteroidota than Firmicutes among the top core nrMAGs (Fig 6B).

**Figure 6:**
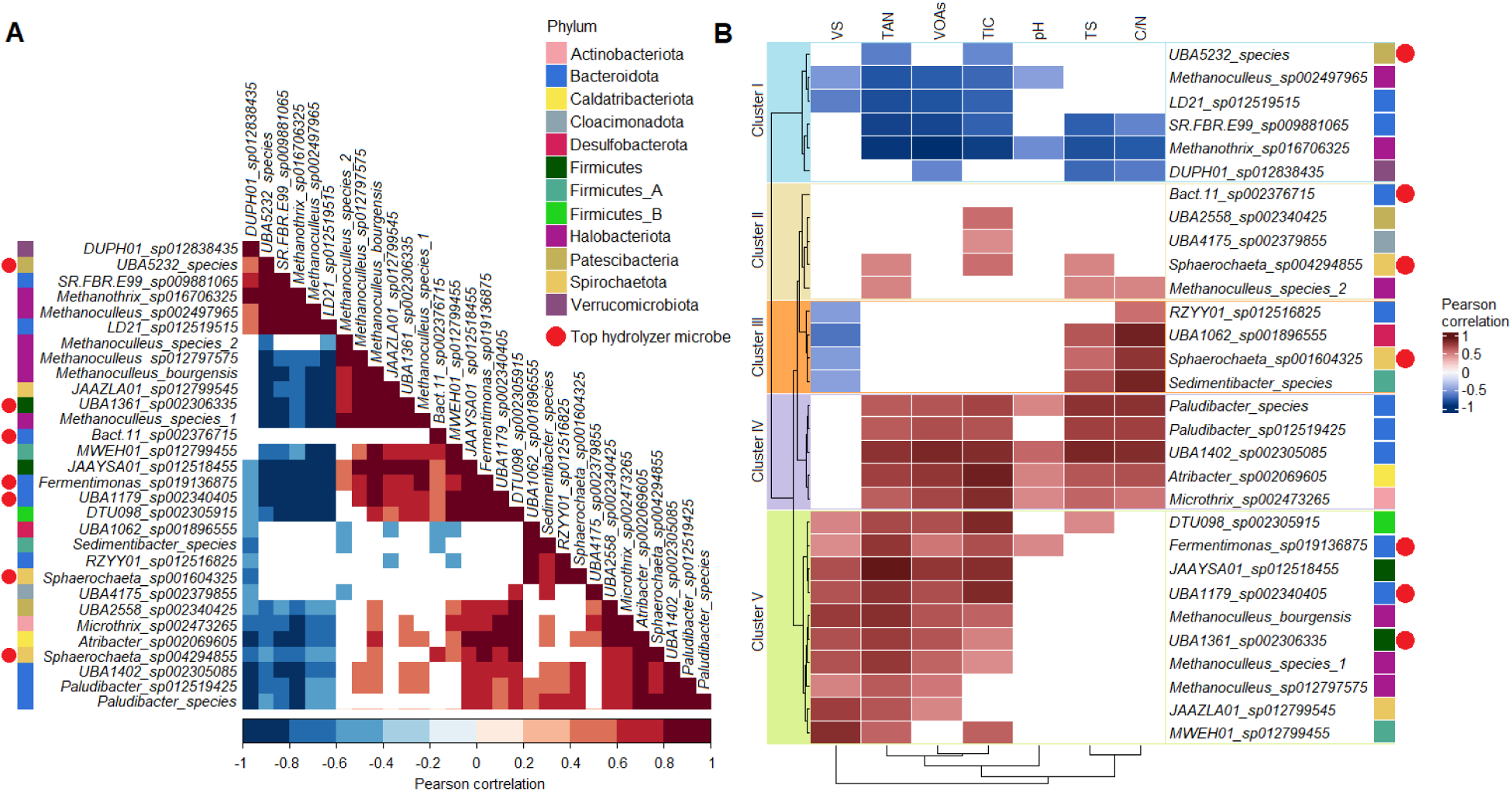
Correlation analysis of the top 30 core nrMAGs. A Pearson correlations between core nrMAGs. The figure shows only significantly positive (>0.5, shown in red) or negative (<−0.5, shown in blue) correlations. VOAs: volatile organic acids, TIC: total inorganic carbon, TAN: total ammonia nitrogen, TS: total solids, VS: volatile solids. B Pearson correlations between core nrMAGs and measured anaerobic digestion parameters. The figure shows only significantly positive (>0.5, shown in red) and negative (<−0.5, shown in blue) correlations. Using the cladogram shown on the left and data for TS, TIC, VOAs and TAN, five clusters were distinguished. These are shown here separated by colour as follows: Cluster I: blue, Cluster II: yellow, Cluster III: orange, Cluster IV: purple and Cluster V: green. Red dots represent the top hydrolysing microbes (n = 7).

The nrMAGs from the genus *Methanoculleus* (phylum Halobacteriota, representing hydrogenotroph methanogens) were primarily found in Cluster V and showed a strong positive correlation with TAN and VOA concentrations. Their abundance was also positively correlated (Pearson’s rho >0.6) with representatives of the phyla Bacteroidota and Firmicutes. It was previously reported that the hydrogenotrophic methanogen *Methanoculleus bourgensis* (belonging to Cluster V in the present study) was positively correlated with TAN [63]. Microorganisms belonging to the phyla Bacteroidota and Firmicutes can produce organic acids, alcohols, hydrogen (H_2_) and carbon dioxide (CO_2_) via acidogenesis and acetogenesis and many of these microbes exist in syntrophy with hydrogenotrophic methanogens. The hydrogenotrophic methanogens consume hydrogen via interspecies hydrogen transfer (IHT) and use the energy gained from reduction of CO_2_ to methane [64]. IHT is a common process in biogas-producing communities and it is thermodynamically feasible to keep the partial pressure of hydrogen low. [12, 65].

Notably, two species of methanogenic archaea, *Methanoculleus sp002497965* and *Methanothrix_sp016706325* (designated MAG_26_2 and MAG_97_2, respectively), were found in Cluster I and showed a negative correlation with microbes belonging to Cluster V. These archaea, which belong to the phylum Halobacteriota, were commonly present in the MWBP microbiome. In contrast, microbes in Cluster V were more prevalent in KBP and SZBP. *Methanoculleus sp002497965* and *Methanothrix_sp016706325* were originally identified in AD microbiomes, but to the best of our knowledge detailed information about these microbes remains unavailable [66–69].

In the present study MAG_26_2 was classified as a strict hydrogenotroph methanogen by metatranscriptomics data. Moreover, it exhibited a negative correlation with TAN concentration (Pearson’s rho = −0.8). Based on genomic data, this archaeon belongs to the class of methanogens capable of maintaining hydrogenotrophic methanogenesis without cytochromes using formate as an electron donor. Typically, formate is oxidised to CO_2_ by formate dehydrogenase, upon which it is further reduced to methane [70]. Here, interspecies formate transfer (IFT) was maintained by microbial partners, namely *SR.FBR.E99_sp002497965* and *LD21_sp012519515* (from the phylum Bacteroidota; MAG_72_3 and MAG_394_2), both of which were positively correlated with MAG_26_2. These MAGs have been previously detected in biogas digesters and were described as protein hydrolysing and amino-acid degrading microbes capable of producing formate; they were also consistently detected alongside formate-utilising methanogens [71].

It is also notable that among the core abundant microbes present we identified *Methanothrix_sp016706325* (MAG_97_2), an archaeon that is apparently capable of both hydrogenotrophic and acetotrophic methanogenesis. Based on its MAG metatranscriptome profile, it primarily performs acetotrophic methanogenesis and can take up electrons via direct interspecies electron transfer (DIET) to drive CO_2_-reducing methanogenesis. Suppl. fig. 1 and Suppl. Table 4 indicates that it possesses highly active Acetyl-CoA and F_420_ biosynthesis pathways as well as active, membrane-associated cytochrome-C biogenesis. This archaeon also showed a positive correlation with the previously mentioned amino-acid degrading microbes, which also have active C-type cytochromes and pili synthesis ability, both of which are essential for DIET (MAG_72_3 and MAG_394_2; Suppl. Table 5) [65, 72–74]. During metabolic processes electrons can be generated and carried by reducing equivalents such as reduced ferredoxin. To re-oxidise these electron carriers, formate production may allow the use of electron disposal routes to conserve energy [71]. Thus, *Methanothrix_sp016706325* (MAG_97_2) may play an essential role in the biogas-producing community by supporting H_2_-independent electron disposal routes [75]. Based on data from the scientific literature, as well as a comprehensive analysis of anaerobic digesters in Danish wastewater treatment plants, representatives of *Methanothrix* showed a negative correlation with both acetate and TAN concentrations [53, 76, 77].

Correlations between microbial community and chemical process parameters implies a distinct electron transfer mechanism during the AD of agricultural biomass and wastewater. These processes are negatively correlated with VOAs and TAN levels and commonly occur when electron disposal routes are abundant. For example, our data and previous studies suggest that protein hydrolysing and amino-acid-degrading microbes –which according to the GTDB taxonomy belong to the phylum Bacteroidota and family VadinHA17 (Suppl. Table 3)– build syntrophic relationships with hydrogenotrophic methanogens to maintain formate production and dispose of electrons via DIET [71].

### 3.6 Top hydrolysers

Carbohydrate-active enzymes (CAZymes) are responsible for the decomposition of polymeric carbohydrates. To characterise this rate limiting step, the metatranscriptome dataset was queried for various CAZymes and signs of CAZyme activity using a combined dataset based on the Pfam and CAZy databases (Fig. 7). Identified CAZymes were then linked to nrMAGs (see section 2.5).

**Figure 7:**
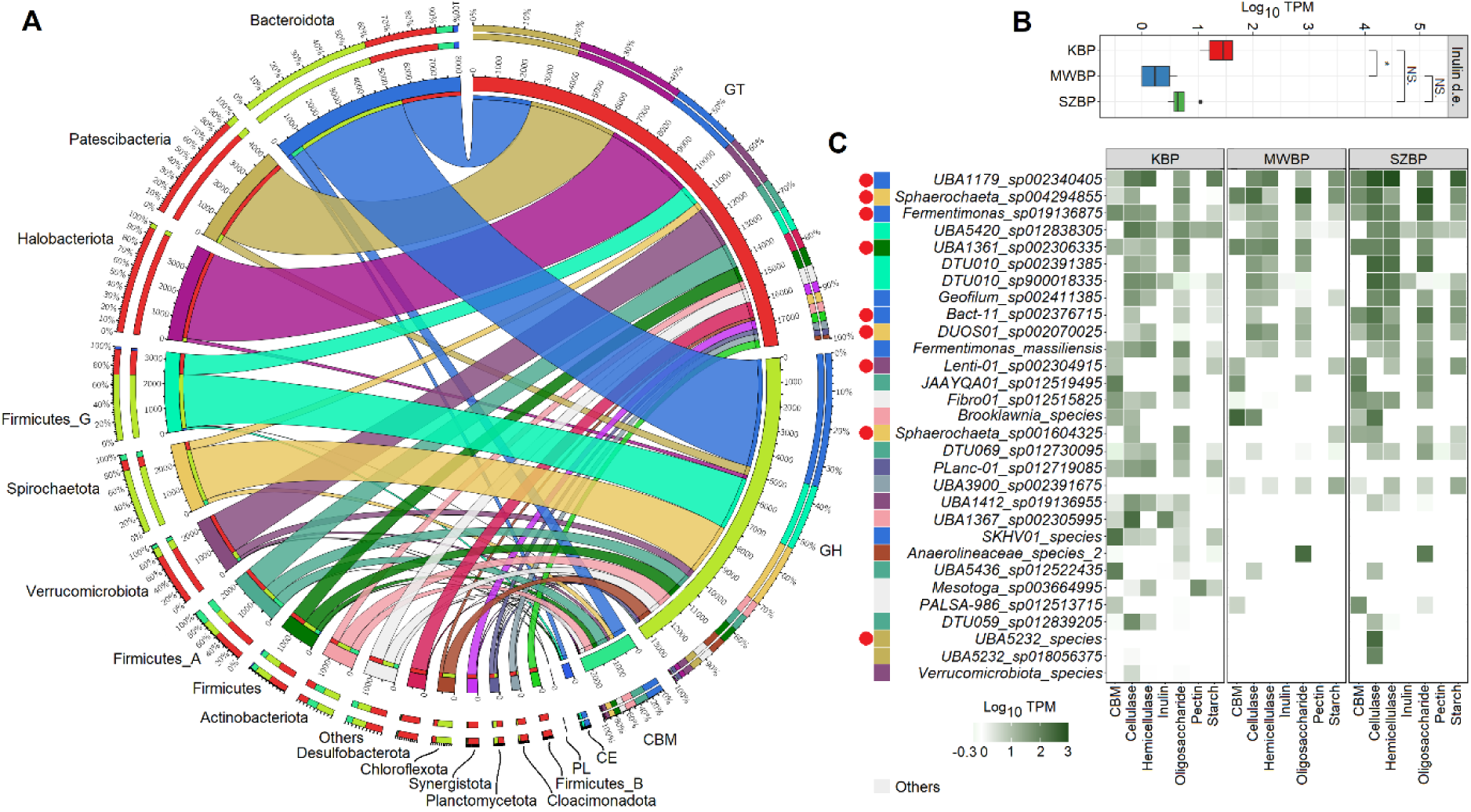
Metatranscriptomic data of identified CAZymes and associated taxa. A Circos plot illustrating identified CAZyme classes and their activity distribution across bacterial phyla. The enzymes were grouped in five CAZyme classes. Glycosyltransferases (GTs) showed the highest activity, followed by glycoside hydrolases (GHs), carbohydrate-binding modules (CBMs), carbohydrate esterases and polysaccharide lyases. Overall, GH and GT activity was detected in all observed microbial phyla. Most (55% TPM) CBM activity was linked to the Firmicutes and Bacteroidota. Box plot presenting significant difference in inulin degradation activity between KBP and the other two BPs (ANOVA: p < 0.05). C Heatmap of the activities of the carbohydrate-binding module (CBM) and glycoside hydrolase (GH) enzyme families for the top 30 microbial species (shown in log10 TPM values). Red dots indicate specific microbes present in the core community (n = 9).

Among the CAZymes identified, the glycoside hydrolases (GHs) are a major enzyme family involved in the degradation of complex carbohydrates (Fig. 7A) [78]. This diverse enzyme family includes cellulases, hemicellulases, pectin, inulin and various oligosaccharide- and starch-degrading enzymes. Here, we detected higher numbers of inulin-degrading enzymes in KBP compared to SZBP and MWBP (ANOVA: p < 0.05) (Fig. 7B). Wheat straw and other terrestrial plant matter contains high levels of inulin, which is used by plants as an energy reserve [79]. According to the metatranscriptome data, hemicellulose-, pectin- and starch-degrading enzymes showed elevated activity in KBP and SZBP (Fig. 7C). However, these differences were not significant (ANOVA: p > 0.05) (Suppl. Fig. 2). Taxonomically, members of the phyla Bacteroidota (representing 35% TPM of GHs) and Firmicutes_G (18% TPM of GHs) were the dominant contributors of GH activity. These phyla have been found to be the main polysaccharide degraders in many biogas-producing communities [62, 80]. Nevertheless, our metatranscriptomics data suggested that members of other phyla such as the Spirochaetota (17% TPM of GHs), Actinobacteriota (5% TPM of GHs) and Chloroflexota (4% TPM of GHs) also expressed complex carbohydrate-degrading enzymes (Fig. 7A) [20, 34]. Interestingly, a combined quantitative assessment of carbohydrate-binding modules (CBM) and GH activity revealed that the top 30 hydrolysing microbes are distributed across multiple phyla (Fig. 7C).

Nine core microbes were detected among highly active hydrolysing nrMAGs. These nine represented about 30% of the total hydrolysers (Fig. 7C). Moreover, seven of the nine core microbes were predominant in the core community (Fig. 7C and Fig. 6A). We found that *UBA5232 species* (MAG_524_1) belongs to Cluster I, *Sphaerochaeta sp001604855* (MAG_59_1) and *Bact-11 sp002376715* (MAG_105_1) to Cluster II, *Sphaerochaeta sp004294325* (MAG_55_1) to Cluster III and *UBA1179 sp002340405*, *UBA1361 sp002306335* and *Fermentimonas sp019136875* (MAG_187_1, _173_1 and _96_1) to Cluster V (Figs. 6 and 7). Members of the top core hydrolysers belonging to Cluster V showed the most widespread interactions. This cluster was directly or indirectly involved via IHT and showed positive associations with *Methanoculleus bourgensis*, *Methanoculleus species1* and *Methanoculleus sp012797575* (Pearson’s rho 0.6–0.9). However, the cluster also showed a negative correlation with *Methanothrix sp016706325* and *Methanoculleus sp002497965* (Pearson’s rho <−0.6). In general, concentrations of TAN, VOAs and TIC were all positively correlated with most active core hydrolysing microorganisms (Pearson’s rho >0.6). However, there were several exceptions involving microbes belonging to Cluster I.

### 3.7 Methanogenesis is the most active pathway in the transcriptome

To further characterise the methane-producing food chain, we performed functional analysis of the nrMAGs by combining metatranscriptome data and data from the KEGG database (Fig. 8). With respect to KEGG classes, we detected no significant differences among the metatranscriptomes of the three biogas plants (ANOVA: p > 0.05) (Fig. 8A). However, the KEGG profiles of the three reactors slightly differed from the profile observed by Ma *et al*. (2021) in their metagenome-based microbial gene catalogue of AD [27] (Fig. 8A). Energy metabolism shows higher activity than carbohydrate metabolism according to our metatranscriptome-based analysis, although the differences between these were not significant (Wilcoxon-test: p > 0.05).

**Figure 8:**
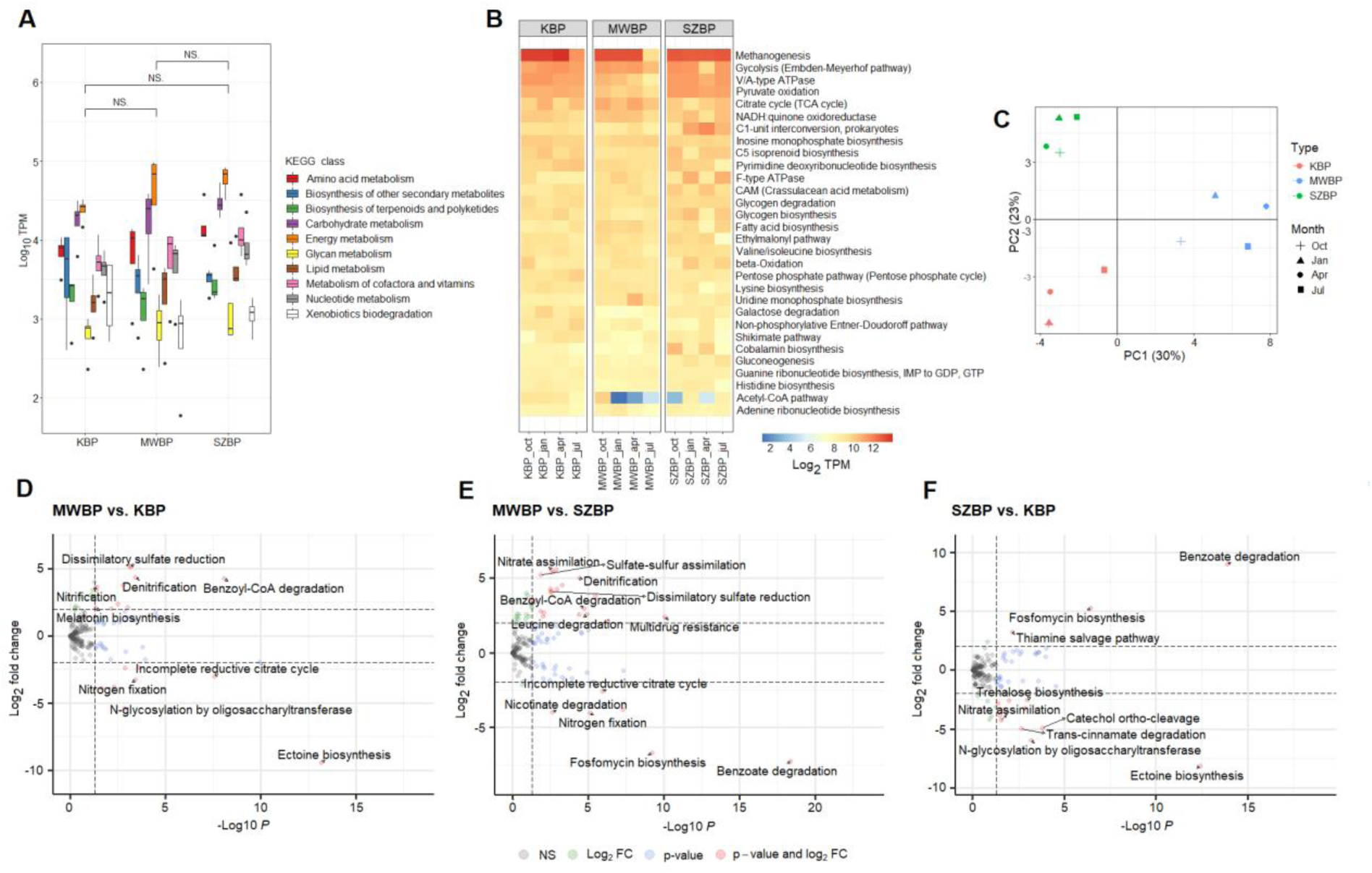
Functional analysis of the three BPs. A Main KEGG classes and their distribution among the biogas plants. An ANOVA detected no significant differences in the levels of KEGG classes among BPs (p > 0.05). B Heatmap of the most active KEGG modules. Shown are the top 30 KEGG modules from all twelve samples based on metatranscriptomic data (shown in TPM). C Principal component analysis of KEGG module functional data. Each symbol is related to a specific BP-sample pair. D, E and F Significantly different KEGG modules between specific BPs. Shown are only those KEGG modules which show the strongest differences. Shown are differences considered to be statistically significant as calculated by Deseq2 (i.e. with p < 0.05 and log2 FC > 2.0).

Further high-resolution analysis of KEGG modules indicated that the methanogenesis pathway was the most active among all main pathways (mean = 14.7% TPM of all KEGG pathways). This finding is remarkable and novel, since metagenome studies have considered methanogenesis to be a ‘rare module’ in the biogas production community [20, 27, 34] (Fig. 8B). It is important to note that while methanogenesis was indeed a rare module based on the metagenome data, the transcriptome data suggested the opposite. Considering this, metatranscriptome analyses, which quantify the biological activities of microbes in complex environments, provide a more accurate representation of microbial life and microbial activity occurring within the community than metagenomics studies.

We identified pyruvate-oxidation, the Embden-Meyerhof pathway and the pentose-phosphate pathway as central carbohydrate metabolism modules. These pathways have been referred to as ‘core modules’ in previous metagenomic investigations [20, 34]. Sugars, coming from hydrolysis, can be converted to pyruvate via the Embden-Meyerhof and pentose-phosphate pathways to produce CO_2_ and electrons. In addition, the acetyl-CoA pathway, fatty acid biosynthesis and beta-oxidation are active among the top 30 pathways related to carbon fixation, as has been previously observed in manure supplemented reactors [20]. This finding was supported by the identification of numerous active modules associated with energy, amino acid and cofactor production, all of which are essential for a well-functioning biogas generation ecosystem. Next, we discovered that KBP and SZBP shared a similar microbial composition and functional profile, while those of MWBP appeared (based on KEGG modules) to be clearly distinct (Fig. 8C). Interestingly, despite functional dissimilarities no differences were detected in the methanogenesis pathway activity of the three BPs (Fig. 8D, E and F). This finding is consistent with biological methane potential measurements (Fig. 1A).

Nevertheless, we also noted modulations in several pathways among the three biogas plant (pCutoff: 0.05, FCcutoff: 2.0) (Fig. 8 D-F). According to our metatranscriptome analysis, these pathways were found to be less abundant and less active, but their occurrence was specific to various biogas plants (reaching to 0.7% TPM of KEGG pathways). The dissimilatory sulfate reduction and denitrification pathways were represented in MWBP (0.3% and 0.2% TPM of KEGG pathways) (Fig. 8D and E). Sulfate-reducing bacteria (e.g. *MWCO01_sp002070115*, MAG_61_2) can reduce sulfate to sulfide by using hydrogen as electron donor for energy conservation. In general, sulfate-reducing bacteria and methanogenic archaea typically coexist in the AD community. They act as competitors for H_2_ and their relationship can be antagonistic or syntrophic, depending on the type of methanogenic archaea (hydrogenotroph, acetotroph, etc.) and the fermentation parameters (sulfate concentration and retention time) [81, 82]. Similarly, in microbes capable of heterotrophic denitrification (e.g. *MWFE01_sp002071445*, MAG_358_2), which may originate from wastewater-sludge substrates [83], their pathways act as a H_2_ sink in the biogas microbiome and may potentially be antagonistic toward hydrogenotroph methanogens [75].

Ectoine biosynthesis and benzoate degradation were identified as unique pathways in KBP and SZBP, representing 0.7% and 0.3% TPM of KEGG pathways, respectively (Fig. 8F). Genome-centric metatranscriptomic data revealed that ectoine is used by *Methanothrix_A_harundinacea_D* (MAG_5_1) as an osmoprotectant, which potentially enhances cell granulation [84]. The benzoate degradation pathway, which requires elevated expression of enoyl-acyl carrier protein reductase can be linked to the novel mixotrophic methanogen *Methanosarcina sp.,* which we found to be common in SZBP (no species-level representative was found in the GTDB and Biogas Microbiome databases; MAG_165_3). This protein is essential for archaea to produce isoprenoid-based lipids, which are membrane-associated molecules that are involved in a wide variety of vital biological functions [85, 86]. The identified multidrug resistance pathway in MWBP was responsible for the increased activity of efflux pump genes, mainly MepA, QacA and MexAB-OprM. These are found in a diverse array of microbes but were specifically detected in MWBP in this study [87] (Fig. 8G). These efflux pumps are not often specific for antibiotics and therefore may confer other evolutionary advantages to their hosts [88].

## 4. Conclusions

In this study, we reconstructed high-quality nrMAGs (non-redundant metagenome-assembled genomes) and conducted a precise metatranscriptome analysis with the assistance of an artificial neural network binning workflow on AD samples taken from industrial-scale biogas reactors. Although the three industrial-scale BPs operated with well-characterised and distinct feedstocks, we observed no differences in their respective methane yields over the observation period. The microbiome composition and the functional repertoires of the KBP and SZBP BPs were more similar to each other than either was to that of MWBP. Multiple bacterial phyla were identified as major hydrolysing microbes. The core microbiome was correlated to the fermentation parameters TAN, VOAs and TIC. This revealed that the interaction between the AD core microbial community and their complex substrate composition is affected by several chemical operational parameters that shape the microbial food chain. Hydrogenotrophic methanogens (Archaea: Halobacteriota) were dominant and positively correlated with the presence of representatives of the bacterial phyla Firmicutes and Bacteroidota, with whom they engage in versatile interspecies transfers. Distinct electron transfer mechanisms used by hydrogenotrophic methanogens in AD have also been found in agricultural biomass and wastewater. A key archaeal species (*Methanothrix_sp016706325*; MAG_97_2) was detected in the core methanogenic community; this species likely plays a key role in shaping the methane-producing microbial strategy. Even though different methane-producing strategies were observed, we did not detect significant differences in the activity of these methanogenesis pathways. This finding was also consistent with our BMP test measurements. Taken together, the results of the present case study confirm that there remain important knowledge gaps in our understanding of the activities and interspecies relationships of members of biogas-producing communities. These can be at least partly resolved via a framework in which genome-resolved metagenome analysis is complemented with a parallel metatranscriptomics approach informed by selective machine-learning algorithms, such as habitat-specific models.

## Data availability

The raw metagenome and metatranscriptome sequences generated and analysed during the current study were deposited in the NCBI SRA under accession number PRJNA929705. High-quality (compl. > 90% cont. < 5%) nrMAGs are deposited in the NCBI SRA under accession numbers from SAMN32989584 to SAMN32989690. The main data generated or analysed for this study are included in this published article and its supplementary information files. Workflows and R scripts are available from the first author on reasonable request.

## Acknowledgements

Roland Wirth designed and performed the bioinformatics analyses and wrote the manuscript. Teur Teur Sally Cheung and Zoltán Bagi performed the experiments and conducted analytical measurements. Prateek Shetty contributed to the meta-omics analyses. Kornél L. Kovács and Gergely Maróti designed the study, wrote the manuscript and discussed the relevant literature. All authors read and approved the final manuscript.

## Competing Interests

The authors declare no competing financial interests.

## Funding

This study has been supported in part by the Hungarian National Research, Development and Innovation Fund (NRDIF) projects: RW, BZ and GM received support from projects PD132145, FK142500, FK123902, FK123899, K143198 and ÚNKP-22-5-SZTE-537. This work was also supported by the Lendület-Programme (GM) and Bolyai Scholarship (WR) of the Hungarian Academy of Sciences (LP2020-5/2020 and BO/00449/22) and by the Széchenyi Plan Plus National Laboratory Programme (National Laboratory for Water Science and Water Security, RRF-2.3.1-21-2022-00008). BZ and KLK received support from 2020-3.1.2-ZFR-KVG-2020-00009 and 2019-2.1.13-TÉT_IN-2020-00016.

## Supplementary content

**Supplementary Table 1:** Summary data of biogas plants. Shown are all main parameters for the studied biogas plants.

**Supplementary Table 2:** Sequencing statistics. Shown are raw and filtered DNA and RNA summary data. Short DNA sequences were assembled by Megahit (see section 2.5). Assembly statistics created by Quast are also shown.

**Supplementary Table 3:** nrMAG statistics. This table shows the main parameters of the reconstructed nrMAGs, including completeness, contamination, genome size and N50. All nrMAGs were taxonomically analysed using the GTDB database (see section 2.5) and the results are summarised in this table. ANI calculation on Biogas Microbiome database the representative MAG code and its taxonomic name were included. Also shown is a novelty analysis based on the GTDB and Biogas Microbiome databases, which includes only those nrMAGs that show both greater than 90% completeness and less than 5% contamination (based on SCGs). MarkerMAG was used to estimate the 16S rRNA gene copy number of MAGs. The final column describes the MAG data improvement compared to GTDB (release 207).

**Supplementary Table 4:** Relative abundances of nrMAGs. Shown are summary data of the relative abundances of nrMAGs expressed in read per kilobase per million (RPKM) values.

**Supplementary Table 5:** Activity of nrMAGs. The MAG taxonomy table contains identified genes found in nrMAGs based on data from the Pfam and KEGG databases. Also shown are the activities of these genes as expressed in TPM (columns 11–22) and read count (columns 23–34). NA refers to hypothetical proteins predicted by Prodigal. The coordinates of the identified genes on the contigs are listed in columns 1–3. Column 10 contains the nrMAG number that the indicated gene belongs to.

**Supplementary Figure 1:** Heat map of the core nrMAGs metatranscriptome. Shown are summary data for the core nrMAGs metatranscriptomics dataset. KEGG modules are listed in order of descending activity. The core microbe names as well as the presence of the top 30 most abundant microbes and the cluster that they belong to are shown at the bottom of the figure. Empty boxes indicate that a specific activity was not found in a specific nrMAG.

**Supplementary Figure 2:** Box plots visualising the calculations of statistically significant differences. CAZymes are summarised according to their families. GH enzyme activities are categorised based on their main activity, (i.e. cellulase, hemicellulase, oligosaccharide-, inulin-, pectin- and starch-degrading enzymes).

## Notes

### Competing Interest Statement

The authors have declared no competing interest.

