## Supplementary figures and images for "Inter-kingdom microbial interactions revealed by a comparative machine-learning guided multi-omics analysis of industrial-scale biogas plants"

### Suppl. Figure 1

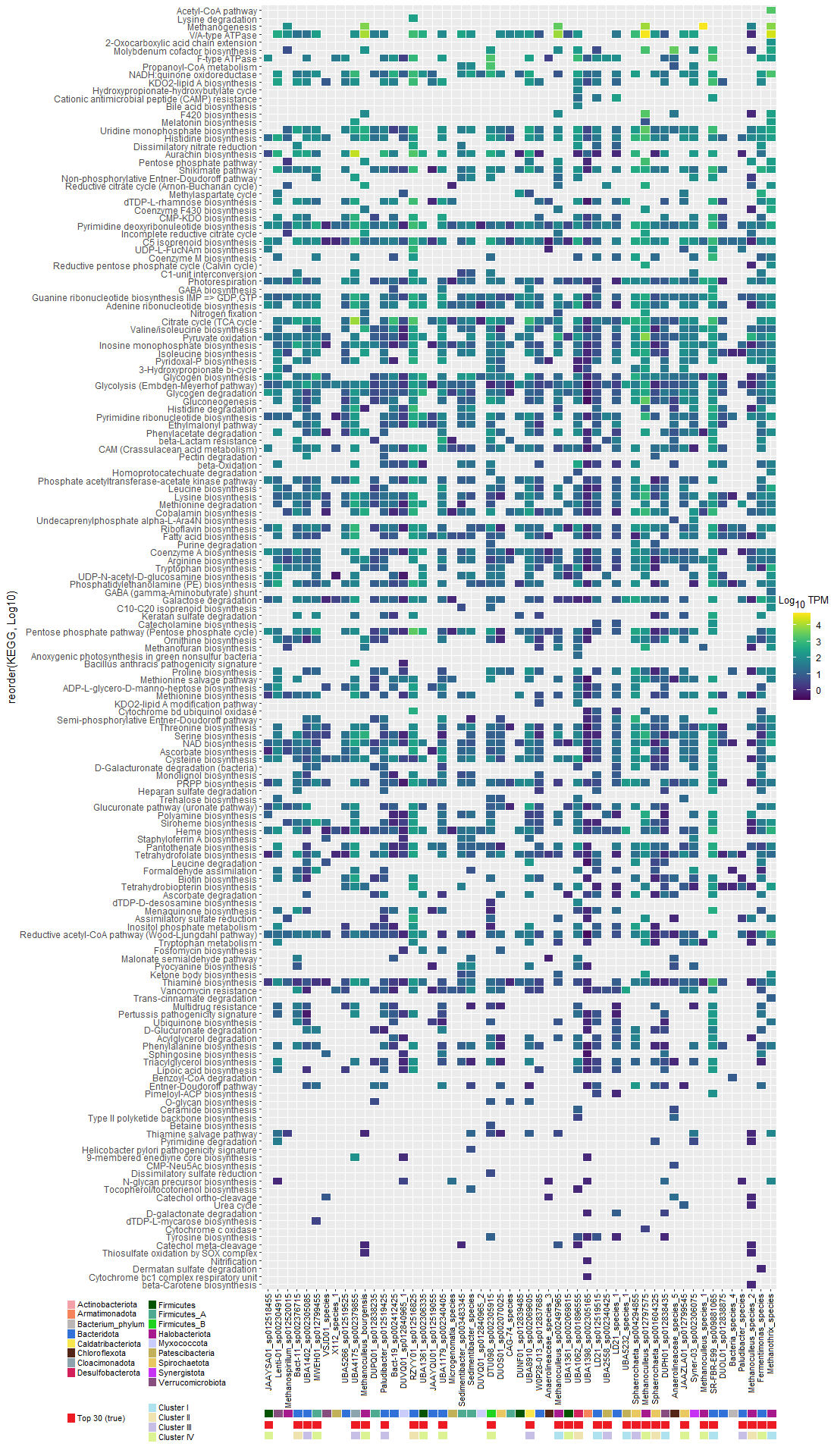

### Suppl. Figure 2

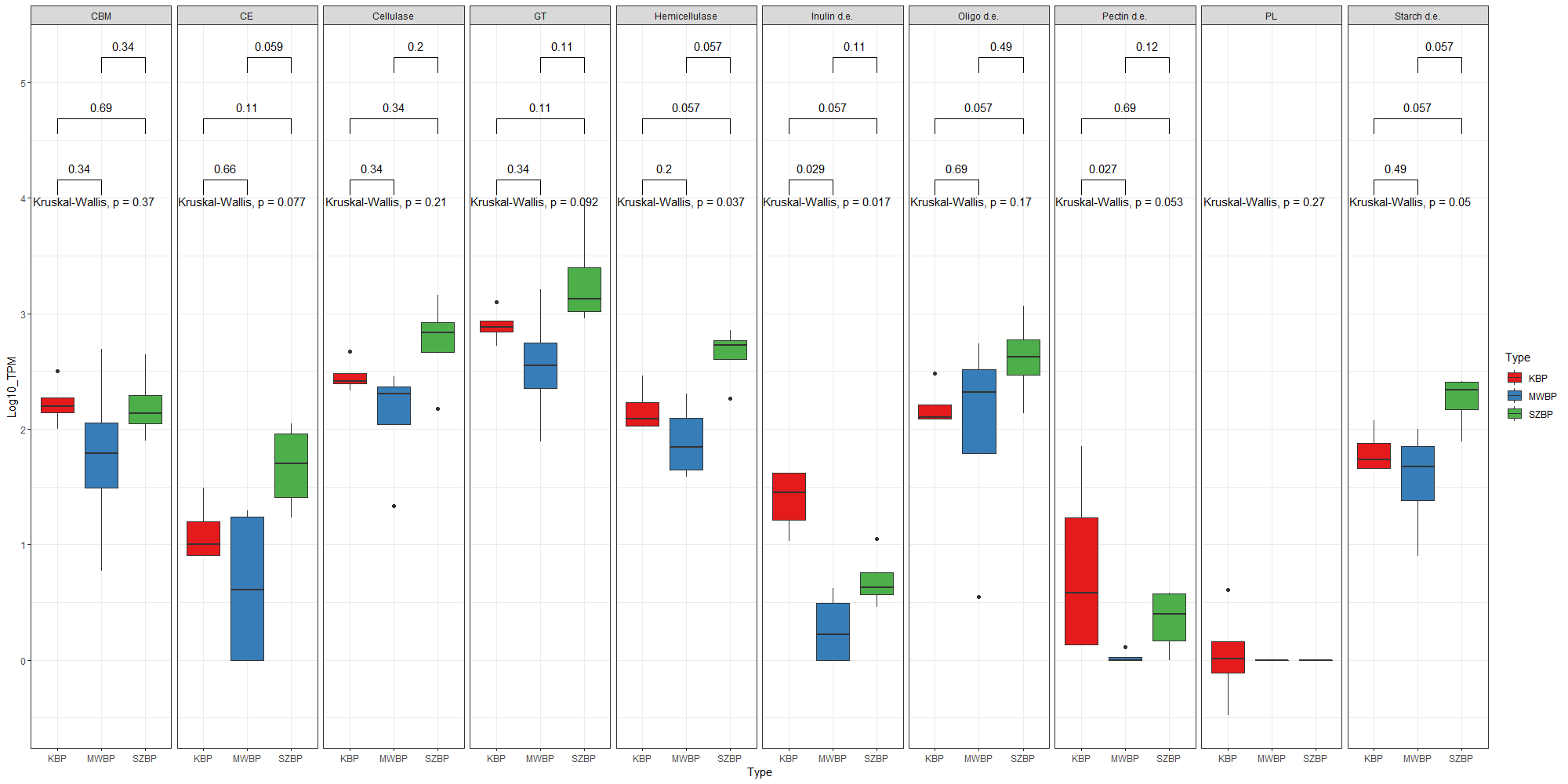
